# Increased levels of HAPLN2, which anchors dense extracellular matrix, in the hippocampus of APOE4 targeted replacement mice

**DOI:** 10.1101/2025.11.09.687435

**Authors:** Samantha Deasy, Matthew Amontree, Zachary Colon, Eric Thorland, Kush Modi, Katie Hummel, Ismary Blanco, Griffin Greco, Kathleen Maguire-Zeiss, Katherine Conant

**Affiliations:** Department of Neurobiology, Duke University School of Medicine, Durham, NC, U.S.A.; Department of Neuroscience, Georgetown University School of Medicine, Washington DC, 20007 U.S.A.; MD/PhD program, Georgetown University School of Medicine, Washington DC, 20007 U.S.A.; Biology Department, Georgetown University College of Arts & Sciences, Washington DC, 20007 U.S.A.; Department of Pharmacology & Physiology, Georgetown University School of Medicine, Washington DC, 20007 U.S.A.; Vanderbilt University, Nashville, TN, U.S.A.

**Keywords:** HAPLN2, APOE, PNN, aggrecan, HTRA1, CSPG sulfation

## Abstract

Hyaluronan and proteoglycan link protein 2 (HAPLN2) / Brain link protein-1 (Bral1) is important for the binding of chondroitin sulfate proteoglycans (CSPGs) to hyaluronan and thus for the formation of specific types of brain extracellular matrix (ECM). It is also significantly increased with aging. Moreover, machine learning has identified it as a brain-derived protein most predictive of Alzheimer’s disease (AD). HAPLN2 binds to CSPGs that may sequester aggregation-prone proteins and also restrict neuronal plasticity. Because the apolipoprotein 4 (APOE4) allele increases AD risk, in the present study we have examined hippocampal lysates from APOE3 and APOE4 targeted replacement (TR) mice using unbiased proteomics, Western blot and hippocampal immunostaining. With proteomics, we observe that HAPLN2 is among the most significantly upregulated proteins in APOE4 mice. Prior work suggests HAPLN2 is particularly important to the assembly of perinodal matrix, and herein we show that it also co-localizes with Wisteria floribunda agglutinin (WFA) positive perineuronal nets (PNNs). PNNs represent a dense form of ECM that can increase GABAergic neurotransmission to alter overall excitatory/inhibitory (E/I) balance and neuronal oscillations important to mood and memory. Proteomics also detected elevated levels of high temperature requirement peptidase-1 (HTRA1), which accumulates in cerebral blood vessels harboring amyloid, in APOE4 mice. In Western blot studies, lysates from APOE4 mice also showed significantly reduced levels chondroitin-6 sulfated proteoglycans, which makes PNNs more susceptible to proteolysis and less inhibitory. In addition, immunostaining studies showed that levels of the PNN component aggrecan were increased in the hippocampus of APOE4 animals. Overall, these findings contribute to an emerging body of literature suggesting that brain extracellular matrix may be altered with aging and other risk factors for AD, and suggest that future studies should assess PNNs, peri-nodal structure and axonal conduction in the background of APOE4.

## Introduction

AD is a disabling and progressive brain disorder that is increasing in prevalence due to an aging world population. In addition to age, risk factors include traumatic brain injury and the relatively common APOE4 allele (15-25% of individuals).

Recent human and animal studies have shown and association between APOE4 and pro-fibrosis molecules including the chemokine CCL5, which promotes ECM deposition in liver and kidney [1-3], and tissue inhibitor of matrix metalloproteases-1 (TIMP-1), which inhibits the activity of ECM degrading proteases [1]. Published data has also linked increased levels of TIMP-1 with brain atrophy [4]. APOE4 is also thought to promote glial activation and brain inflammation [5], which in the long term can enhance ECM deposition [6].

Increases in brain ECM could contribute to AD through varied mechanisms. Increases in perineuronal net density can impair cognition [7-9], and peri-nodal ECM deposition can impair axonal conduction [10]. And though ECM rich structures can also be protective [11], select ECM components could potentially sequester β- [12, 13]. Amyloid sequestration can also occur in brain regions including the blood brain barrier, and thus contribute to BBB injury with anti-amyloid antibodies [14].

Increases in brain ECM may also disturb neuronal network dynamics. Of interest, human APOE4 expressing mice have reduced gamma oscillation power as compared to their human APOE3 expressing counterparts [15], and it is tempting to speculate that this could be in some part due to an increase in ECM deposition. Gamma oscillation power, which is generally reduced in AD patients [16, 17], is increased in rodents and *ex vivo* murine slices with PNN attenuation [8, 18, 19]. PNNs are predominantly localized to fast spiking parvalbumin (PV) positive interneurons, and facilitate their firing through multiple mechanisms including reduced membrane capacitance [20]. Excess PNN deposition may thus increase interneuron activity and in turn reduce excitatory neurotransmission [21].

To better understand whether specific ECM components are altered in association with APOE4, we performed unbiased proteomics on hippocampal lysates from APOE3 and APOE4 targeted replacement mice. In addition, we complemented these studies with targeted analysis of select ECM components by Western blot and immunochemical staining.

## Materials and Methods

### Subjects

The present study used aged and sex matched groups of human APOE3 and APOE4 targeted replacement mice for proteomics/Western blot and, due to limited availability of the TR mice, age and sex matched groups of human APOE3 and APOE4 knock-in mice for immunostaining. Previous studies have shown that the two human APOE3 and APOE4 models show similar biochemical changes [22]. 12 month old mice were used for proteomics and western blot and included 3 females and 7 males per group. 12-14 month old mice were used for immunostaining and included females only. Both strains are on a C57/bl6 background and further details on their origin are included in prior publications [22]. Because mouse APOE has different properties and low homology with human isoforms, wild type mice do not represent an ideal model for studying differences with respect to human specific isoforms and thus published studies have instead specifically compared the humanized APOE alleles [1, 23].

### Care of mice

Mice were group housed (4–5 mice/cage), unless otherwise specified, in a temperature and humidity-controlled facility at Georgetown’s Department of Comparative Medicine maintained on a 12-hour light cycle (06:00–18:00). Mice had ad libitum access to food and water. Mice were monitored daily for overall health. All efforts were made to ease pain and distress and minimize the number of mice required. All procedures were performed in accordance with the Institutional Animal Care and Use Committee of Georgetown University, Washington DC, USA (Protocol # 2016-1117), and consistent with the Guide for the Care and Use of Laboratory Animals. Prior to euthanasia by decapitation, mice were deeply anaesthetized with inhaled isoflurane.

### Preparation of Brain lysates

Following anesthesia with isoflurane and euthanasia via decapitation, hippocampi were micro-dissected. Lysates were placed in a microcentrifuge tube containing Radio Immunoprecipitation (RIPA) buffer [50 mM Tris, pH 7.5, 150 mM NaCl, 0.1% sodium dodecyl sulfate, 1% octylphenoxypoly (ethyleneoxy) ethanol, branched, and 1× protease and phosphatase cocktail (Thermo Scientific). Lysates were sonicated for 10 seconds, cooled on ice, and centrifuged for 15 minutes at 13,300 X g at 4 °C. Lysate supernatants were saved for protein analyses.

### Proteomics

Proteomics was performed using methods as described in a previous publication from the lab [24]. Briefly, an equal amount of protein from each lysate (100 μg) was processed for proteomics according to a recently reported procedure [25]. Samples were suspended with 50 μl of 5%SDS buffer (containing 50□mM TEABC and 20□mM DTT) and heated for 10 minutes at 95°C. After cooling down to room temperature, 4 μl of 500□mM iodoacetamide in 5% SDS solution was added to a final concentration of 40□mM and incubated in the dark for 30Lminutes. Undissolved matter was centrifuged for 8 minutes at 13,000 x g. The supernatant was saved and used for downstream processing using a S-Trap column (ProtiFi, LLC). Proteins were digested with sequencing-grade Lys-C/trypsin (Promega) by incubation at 37°C overnight. The resulting peptides were eluted and dried down with a SpeedVac (Fisher Scientific) prior to mass spec analysis with a nanoAcquity UPLC system (Waters) coupled with Orbitrap Fusion Lumos mass spectrometer (Thermo Fisher). Analysis of DIA raw files was done using Spectronaut (Biognosys, v17) with a hybrid library. All settings were default. In brief, dynamic retention time prediction with local regression calibration was selected. Interference correction on MS and MS2 levels was enabled. The Q-value Cutoff was set to 1% at peptide precursor and protein levels using scrambled decoy generation and dynamic size at 0.1 fractions of library size. Proteomics analyses were performed separately on two cohorts of 10 mice within the group of 20 animals used for Western blot. The first proteomics cohort compared two groups of 2 male and 3 female APOE3 with 2 male and 3 female APOE4 animals, and the second cohort compared 5 male APOE3 with 5 male APOE4 animals. Proteomics results shown in column graphs represent the within group normalized data which was then combined.

### ELISA

Hyaluronic Acid (HA) in APOE-TR mice was measured using the Hyaluronan DuoSet ELISA (R&D Systems, Cat#DY3614). HA values were normalized to the total protein concentration.

### Western blot

Total protein was quantified using the Pierce BCA kit (ThermoFisher, Cat#). For immunoprobing of CSPGs, samples were digested with Chondroitinase ABC (ChABC) (Sigma-Aldrich, Cat# C3667) for 30 minutes at 37°C. For immunoprobing of HAPLNs, samples did not undergo ChABC treatment and were directly mixed with 4x Laemmli buffer + beta-mercaptoethanol, heated at 85° C for 10 minutes, cooled on ice, and then loaded on hand-casted single-percentage SDS-PAGE gels (6 and 12%). The gels were transferred to nitrocellulose membranes (Trans-Blot Turbo Transfer, Bio-Rad, catalog #1704159), stained with ponceau S, and blocked for 1 hour in 5% nonfat milk blotting-Grade buffer (Bio-Rad, Cat# 1706404). All primary antibodies were diluted in 3% bovine serum albumin (BSA)L+L0.05% tween-20 (v/v in TBS), and membranes were incubated at 4°C on orbital shaker overnight. The next day, membranes were washed 3 times for 5–10□minutes each with 0.05% tween-20 (v/v in TBS) and then incubated for 1□hour in 5% nonfat milk blotting-Grade buffer +0.05% tween-20 (v/v in TBS, Bio-Rad, Cat# 1706404) with the appropriate secondary HRP-conjugated antibody at 1:5000 dilution at room temperature on an orbital shaker. Membranes were subsequently washed three times for 5–10□minutes each with 0.05% tween (v/v in TBS) and twice with TBS (no detergent) prior to chemiluminescent detection with SuperSignal™ West Pico PLUS Chemiluminescent Substrate (ThermoFisher, Cat# 34579) or SuperSignal™ West Femto Maximum Sensitivity Substrate (ThermoFisher, Cat#34095). The West Femto substrate was diluted 1:6 in West Pico PLUS when West Pico PLUS demonstrated insufficient chemiluminescent detection. Western blots were stripped with restore western blotting stripping buffer (Thermo Scientific™, Cat# 21059) for 30□min at room temperature on orbital shaker, washed three times for 5–10□minutes with 0.05% tween (v/v in TBS), and probed with subsequent primary antibody. All primary antibodies were initially tested on fresh western blots to ensure lack of cross-reactivity that may occur with stripping and re-probing membranes. The following primary antibodies were used: 6-Chondrotin sulfate (1:100, AMSBIO, Cat# 270433-CS), HAPLN1 (1:2000, NSJ BioReagent, Cat# RQ5347), HAPLN2 (1:2000, Antibodies inc, Cat# 75-341). The secondary antibodies used are the following: Goat anti-Rabbit IgG (H□+□□)-HRP (1:5000, Invitrogen, Cat# 31460) and Goat anti-Mouse IgG (H□+□□)-HRP (1:5000, Invitrogen, Cat# 31430).

### Immunochemical staining and analyses

Mice were euthanatized, brains were collected and stored in 4% paraformaldehyde at 4 °C for 24 hours. The next day brains were washed once with PBS and then stored in 30% sucrose (w/v in PBS) at 4 °C. The microtome was used to obtain 30 µM slices and slices were stored in cryoprotectant at -20 °C. <u>For the aggrecan and WFA immunostaining in figure 6:</u> n=4-5, 12-14 month old human APOE3-KI and APOE4-KI mice were used and approximately 12 hippocampal hemi-slices per animal were quantified for aggrecan and 10-11 for WFA. For figure 2, n=4 12-14 month human APOE3-KI and APOE4-KI mice were used and approximately 3-4 hippocampal hemi-slices per animal were quantified for HAPLN2 and WFA.A blinded investigator outlined PNN-like structures and used Image-J software to get the mean intensity value for each. The following primary and secondary antibodies were used for immunostaining in figures 2 and 6: HAPLN2: (Antibodies Inc, Cat# N364/10), HAPLN1: (NSJ Bioregents, Cat# RQ5347), Aggrecan (EMD Millipore, Cat# AB1031), WFA (Vector laboratories, 488 Cat# FL-1351)1351, Goat Anti-Rabbit secondary: Invitrogen (Alexafluor 633 IgG), Goat Anti-Mouse secondary, Invitrogen, (Alexafluor 555 IgG).

**Figure 1.**
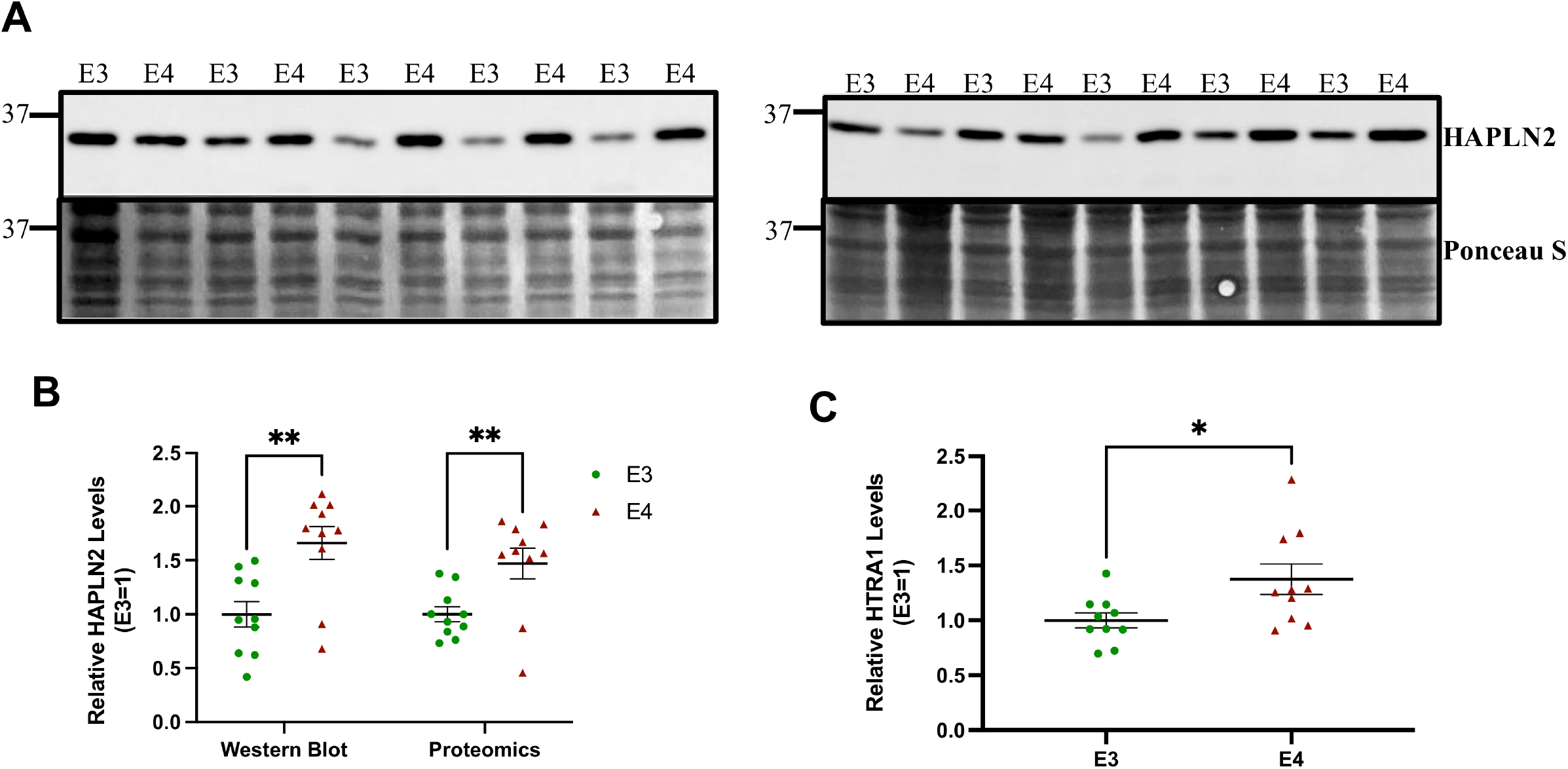
HAPLN2 and HTRA1 levels are increased in APOE4/4TR mice. A) Western blot for HAPLN2 in APOE TR hippocampal lysates. Quantification of Western blot and proteomic HAPLN2 levels in APOE TR mice are shown in B, and quantification of HTRA1 levels (proteomics) in APOE-TR mice is shown in C. Graphs display the mean ± SEM, n_E3_= 10, n_E4_= 10, and significance was tested with Students unpaired two-tailed T test, *p< 0.05, **p<0.01. Parameters for the HAPLN2 Western were: [t(18)= 3.420, p=0.0031], for HAPLN2 proteomics parameters were [t(18)= 2.951, p=0.0085], and for HTRA1 proteomics parameters were [t(18)= 2.422, p=0.0262].

**Figure 2.**
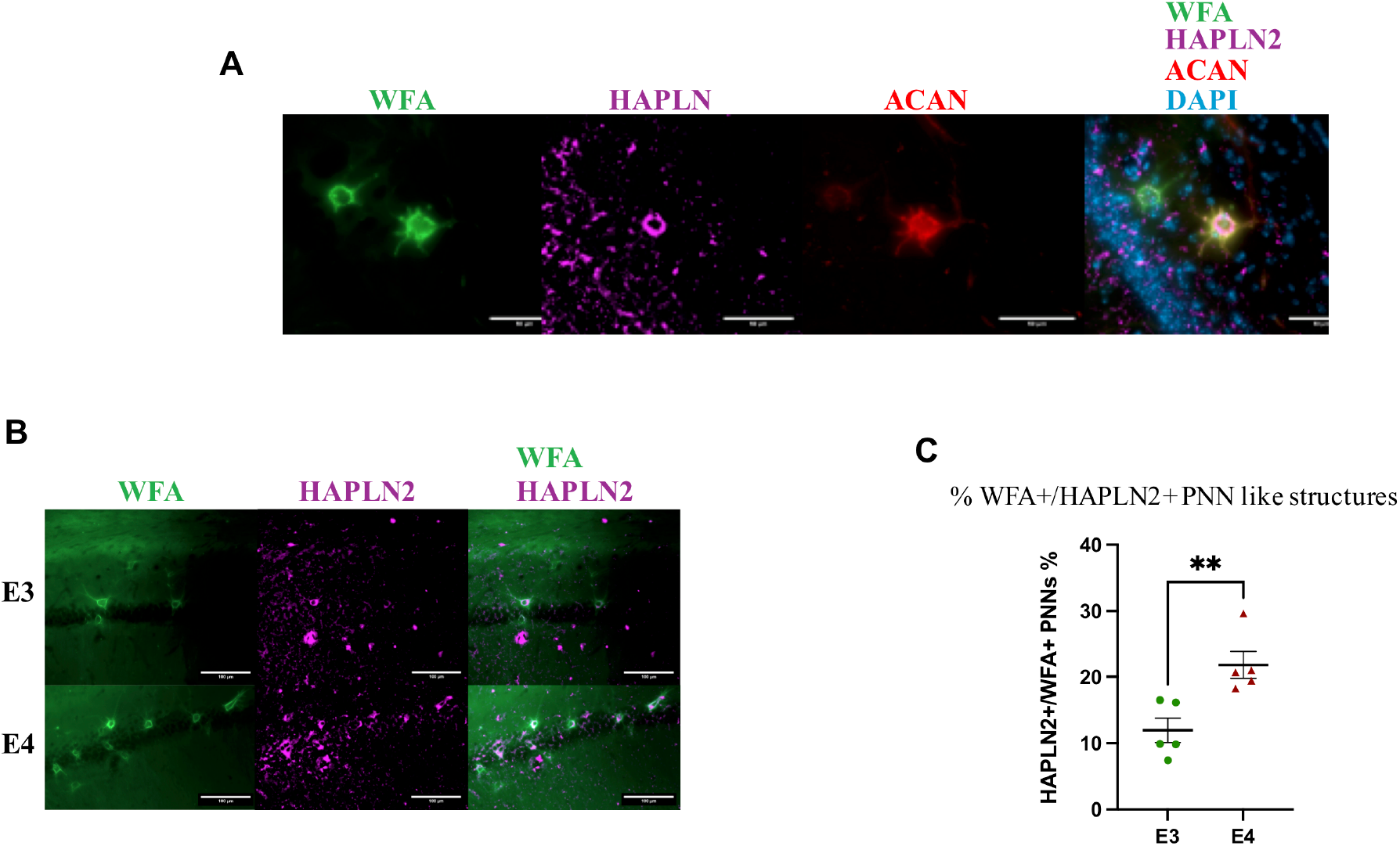
HAPLN2 immunoreactivity colocalizes with that for ACAN and WFA. Immuno-stained hippocampal sections show expression of WFA, HAPLN2, and WFA in PNN-like structures. As shown in Figure 2A, in which HAPLN2 immunostaining is shown in magenta and WFA in green. HAPLN2 and ACAN can be seen in 1 of 2 WFA positive PNN like structures in the image. There is also punctate-like HAPLN2 staining which may be localized to loose ECM and represent aggregates. Shown in Figure 2B is a lower power image of hippocampal staining for HALPN2 (magenta) and the PNN marker WFA (green) in representative APOE3 and APOE4 hippocampal sections. Consistent with quantified data in Figure 2C, the percent WFA+/HAPLN2+ nets are increased in APOE4 hippocampi.

### Statistics

Student’s Two-tailed T-test was used in densiometric analysis comparing protein levels between APOE3-TR and APOE4-TR mice. Two-way analysis of covariance (ANOVA) was used to compare aggrecan fluorescent intensity in APOE3-KI and APOE4-KI and between hippocampal subregions. Outlier analysis was performed using Grub’s testing (alpha=0.05) All statistical analyses were performed in GraphPad Prism version 9.0.

## Results

### I. Proteomics identifies HAPLN2 among the most significantly upregulated proteins in APOE4 mice

Hyaluronan and proteoglycan link protein 2 (HAPLN2) binds to CSPGs that are localized to PNNs as well as peri-nodal sites along CNS axons/at nodes of Ranvier. HAPLN2 is highly expressed in the hippocampus and HAPLN2 can associate with PNN components including versican, brevican and tenascin [26, 27] . Moreover, HAPLN2 levels are increased with aging and HAPLN2 forms aggregates and may also promote the aggregation of α-synuclein [28, 29]. As shown in Figure 1A, proteomics analyses demonstrate that HAPLN2 levels are increased in human APOE4 expressing mice. Western blot analyses also show increased HAPLN2 in APOE4 murine brain lysates (Figure 1B). Differences in HAPLN1 were not observed (supplemental Figure 1). Proteomics also detected elevated levels of high temperature related protein A1 (HTRA1) in hippocampal lysates from APOE4 mice (Figure 1C). This is a serine protease that has been implicated in cerebral small vessel disease and has been shown to colocalize with vascular amyloid [30].

### II. HAPLN2 colocalizes with WFA and HAPLN1 positive PNNs

HAPLN1 and HAPLN4 have been shown to co-localize with PNNs, and HAPLN1 knockout mice show significantly attenuated, though not absent, PNNs [31]. HAPLN2 can also bind to PNN components, and in recent work, HAPLN2 has been shown to co-localize with PNN-like structures in the enteric nervous system [32]. To address the possibility HAPLN2 might also colocalize with PNNs in the hippocampus, and thus potentially increase brain PNN levels if its expression were pathologically increased, we immuno-stained hippocampal sections for WFA, ACAN and HAPLN2. As shown in Figure 2A, in which HAPLN2 immunostaining is shown in magenta and WFA in green, HAPLN2 and ACAN can be seen in 1 of 2 WFA positive PNN like structures in the image. There is also punctate-like HAPLN2 staining which may be localized to loose ECM and represent aggregates. Shown in Figure 2B is a lower power image of hippocampal staining for HALPN2 (magenta) and the PNN marker WFA (green) in representative APOE3 and APOE4 hippocampal sections. Consistent with quantified data in Figure 2C, the percent WFA+/HAPLN2+ nets are increased in APOE4 hippocampi.

### III. HAPLN1 levels do not differ as a function of APOE4

Because HAPLN1 deficiency can significantly attenuate PNNs, we also examined HAPLN1 levels in APOE3 and APOE4 TR mice. The relative levels detected by proteomics and Western blot are shown in Figure 3. As can be appreciated from this data, HAPLN1 levels did not differ as a function of APOE4 genotype [t(18)= 0.134, p=0.895].

**Figure 3.**
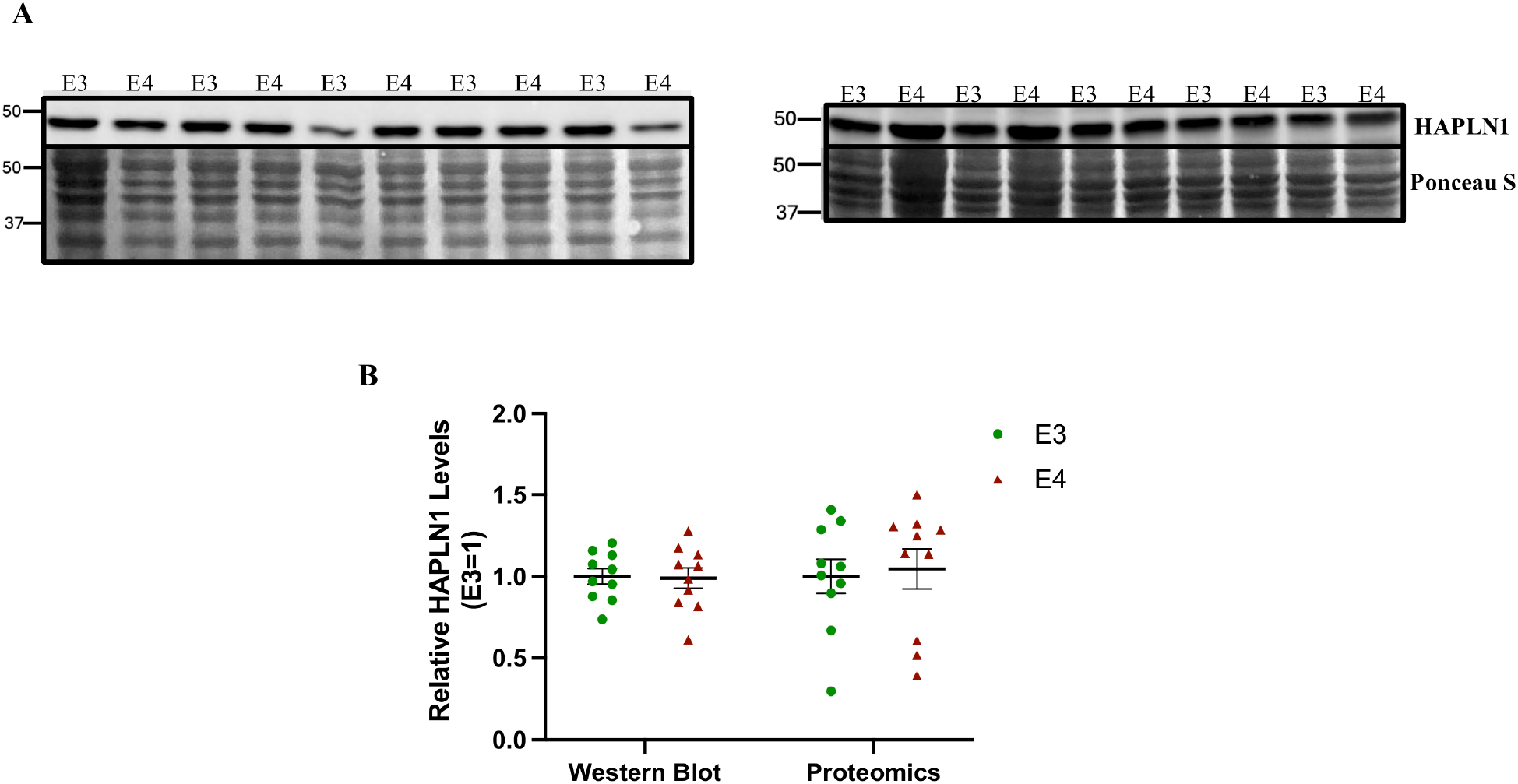
HAPLN1 levels do not differ as a function of APOE4. A) Western blot for HAPLN1 in APOE TR hippocampal lysates. B) Quantification of Western blot and proteomic HAPLN1 levels in APOE TR mice, Graphs display the mean ± SEM, n_E3_= 10, n_E4_= 10, and significance was tested with Students unpaired two-tailed T test. HAPLN1 Western blots shown in A were first probed with HAPLN2, stripped, and then subsequently probed with HAPLN1 antibody.

### IV. Chondroitin-6 sulfated CSPGs, which make PNNs less inhibitory, are reduced in APOE4 mice

Site specific sulfation of chondroitin sulfates allows for selective interactions with different molecules and confers differences in CSPG function [33]. Chondroitin sulfate is composed of a D-glucuronic acid and N-acetylgalactosomine with sulfation potentially occurring at C2 of the former and C4 or C6 of the latter. C6 sulfated CSPGs can make PNNs more sensitive to proteolysis and thus less inhibitory. C4 CSPGs increase while C6 CSPGs decrease during the critical period of plasticity [34]. C6 CSPGs are also reduced with aging [35], and overexpression of the C6 targeting sulfotransferase improves memory in aged mice [36].

As shown in Figure 4, hippocampal homogenates from APOE4 mice show reduced levels of C6 CSPGs as determined by Western blot. APOE4 associated changes in CSPG sulfation are consistent with previous work suggesting that APOE4 can be associated with an accelerated aging phenotype. It is also of interest that changes in the sulfation of CSPGs may occur in AD and that C4 CSPGs can also be targeted with an antibody to improve cognition in a mouse model of tauopathy [37, 38].

**Figure 4.**
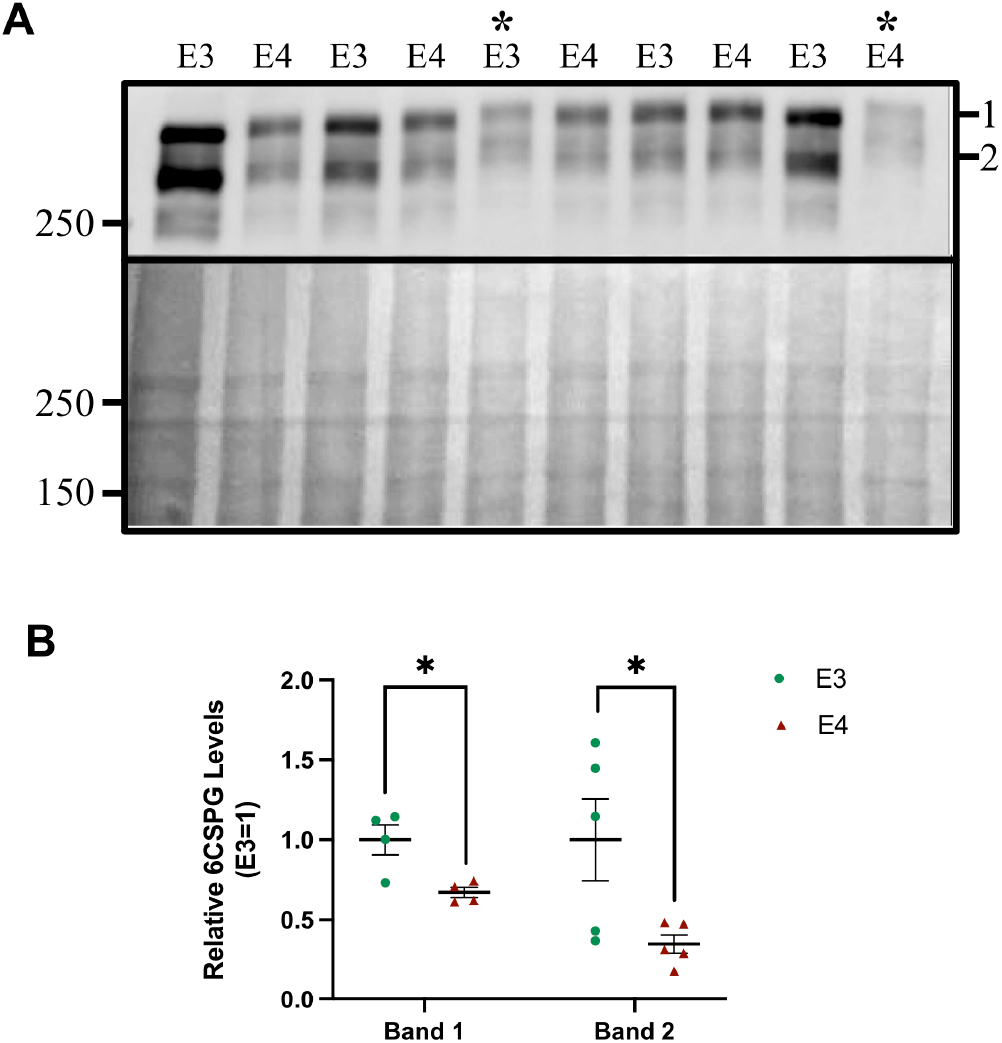
High molecular weight 6CSPGs are decreased in APOE4 TR mice. A) Western blot showing two high MW 6CSPG bands labeled “1” and “2”. B) Densitometry of bands “1” and “2”. Two values were identified as outliers via Grub’s test (alpha=0.05) for band 1 and are highlighted on the western blot with an asterisk. Shown is mean ± SEM, with statistical analysis via Student’s unpaired two-tailed T test (*p<0.05). For both band 1 (n_E3_= 4, n_E4_= 4) and band 2 (n_E3_= 5, n_E4_= 5) there was a significant difference in the relative levels of 6CSPG (band 1 [t(6)= 3.283, p=0.0168]), (band 2 [t(8)= 2.481, p=0.0381]) .

### V. Hyaluronan levels do not differ as a function of APOE4

Recent work has shown that hyaluronan, which is an important component of loose ECM in addition to PNNs, is increased with aging [39]. It is also increased in the setting of microglial depletion [40]. We therefore measured hyaluronan levels in brain lysates from APOE3 and APOE4 TR mice (Figure 5). In contrast to HAPLN2 and C6 CSPGs which have been shown to change with age, and as shown herein with APOE4, hyaluronan levels did not differ as a function of APOE4 [t(16)=1.238, p=0.2337].

**Figure 5.**
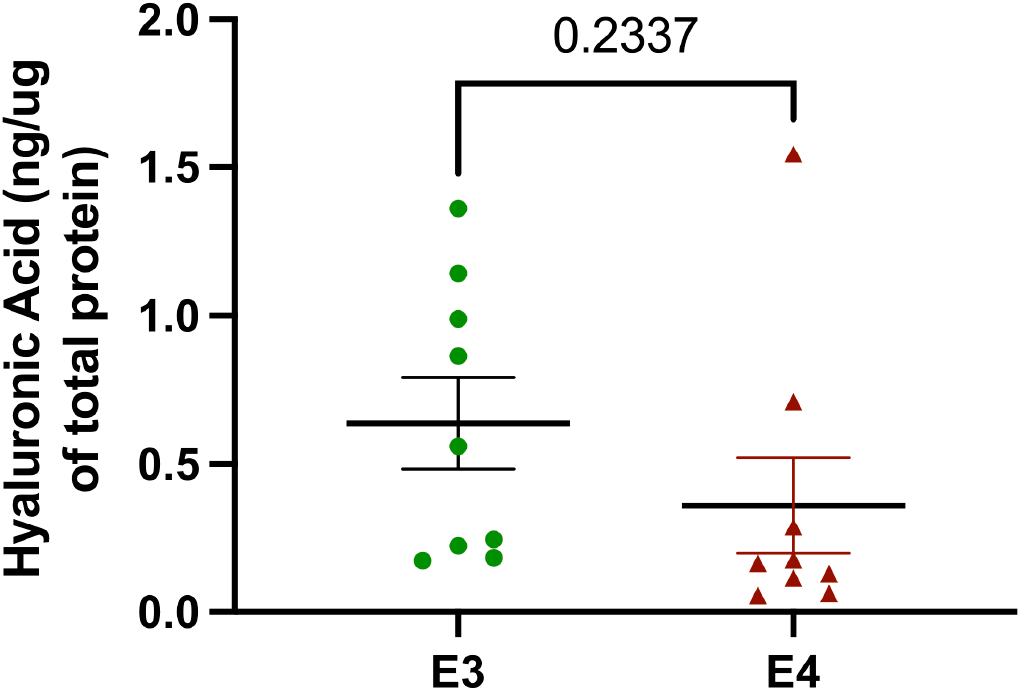
Hyaluronan levels do not differ as a function of APOE4. Quantification of hyaluronic acid ELISA. Two values were identified as outliers via Grub’s test (one outlier in APOE3-TR and one outlier in APOE4-TR). Shown is mean ± SEM, n_E3_= 9, n_E4_= 9, with statistical analysis via Student’s unpaired two-tailed T test.

**Figure 6.**
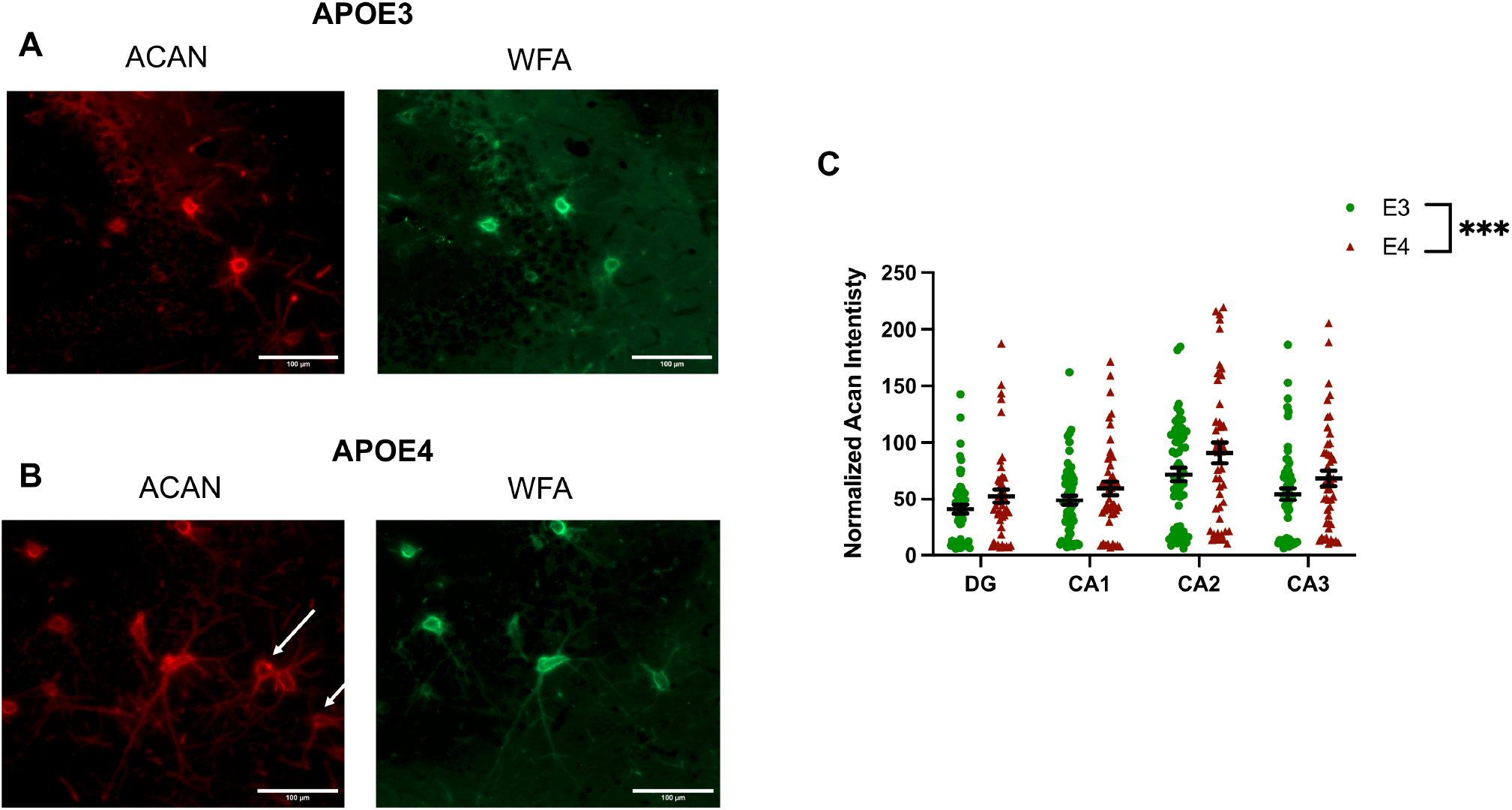
Hippocampal aggrecan staining intensity is increased in APOE4 expressing mice. (A & B) are representative aggrecan (ACAN) (red) and WFA (green) immunostaining images from CA3 hippocampus of female APOE3 (A) and APOE4 (B) murine hippocampi as indicated. The scale bars represent 100 μm. Some cells (white arrows) are ACAN positive but WFA negative. Shown in C) are ACAN intensity measures for hippocampal PNN like structures. Each dot represents the average aggrecan intensity for each visible PNN like structure (proximal dendrites were not included) within the hippocampal region indicated within each hemi-hippocampal slice. Approximately 12 hemi-slices per animal were imaged for 4-5 animals per genotype. The 2-way ANOVA for region and genotype showed a significant genotype difference for the whole hippocampus (*p*= 0.001)

### VI. Aggrecan staining intensity is increased in the hippocampus of APOE4 expressing mice

While WFA is commonly used to stain PNNs, it preferentially recognizes and binds the non-sulfated chondroitin isomer which differs between brain regions. Aggrecan is a PNN component that is highly expressed in the hippocampus and often used in combination or in lieu of WFA [41]. There is also an increased abundance of WFA-negative, aggrecan-positive PNNs in hippocampus [42], and aggrecan is more likely to be localized to PV neurons than are other PNN components such as brevican [43].

As shown in Figure 6, hippocampal aggrecan intensity was increased in APOE4 compared to APOE3 female mice. Shown is the mean ± SEM from a two-way ANOVA (***p<0.001).

## Discussion

Accumulating evidence suggests that excess ECM or PNN accumulation can occur with conditions in which chronic stress and/or inflammation play a role [6]. For example, increased depressive like behavior and increased PNN levels have been described in mice following repeated social defeat stress or chronic treatment with corticosterone [8, 44]. Chronic stress or inflammation can increase ECM remodeling, and in the long term this favors increased matrix deposition. This process is also observed in sites beyond the brain and may underlie pathology observed in pulmonary, renal and liver fibrosis [3, 45].

In the current study, we examined the possibility that mice expressing human APOE4, which is associated with a pro-inflammatory brain state [2, 5], would show increases in specific ECM and/or PNN components. The possibility that APOE4 expression could engender increases in ECM components is also based on prior work from our laboratory and others showing that the APOE4 allele is linked to an increase in varied effectors of ECM deposition including chemokines and tissue inhibitors of metalloproteinases [1, 2].

As shown herein, we find that APOE4 targeted replacement mice show increased levels of HAPLN2, which binds ECM proteins including versican and brevican [46, 47]. While HAPLN1 deficient mice show attenuated PNNs [31], suggesting HAPLN1 is an important lynchpin for PNNs, as shown herein we observe that HAPLN2 may also colocalize with PNNs. One possibility is that with aging or other conditions that increase HAPLN2, its relative contribution to PNNs could increase. Consistent with an aging related increase in HAPLN2 is a recent study that showed HAPLN2 increases in extracellular vesicles from aged mice [48]. Of interest, this study also showed that aged mice showed increased immunostaining for aggrecan, but not WFA, in several brain regions. Further studies will however be necessary to determine whether PNN associated HAPLN2 is increased in the background of APOE4. Further studies should also examine HAPLN2 at nodes of Ranvier, where it is known to be expressed and essential for salutatory conduction [49-51]. In select circumstances excess or pathologically increased perinodal ECM might impair axon conduction [10], and in one study using a murine model of AD, a deficit in axonal conduction has been observed [52]. Recent work has also linked APOE4 to impaired cholesterol localization and reduced myelination, which could also be associated with deficits in axonal conduction [53].

In the present study we also observe increased HTRA1 in the hippocampi of APOE4 mice. HTRA1 is an astrocyte derived protease that has been found to accumulate with cerebral vessels harboring amyloid in a rat model of cerebral angiopathy [30]. It may also be dysregulated in major depressive disorder and murine models of the same [54]. It is thus tempting to speculate that HTRA1 may contribute to increased matrix remodeling in the setting of chronic stress and/or inflammation. Future studies should also examine the potential for HTRA1 to contribute to increased blood brain barrier remodeling that increases the risk of micro-bleeds in the background of APOE4. Additional significantly changed proteins in the proteomics analyses included 0610012G03Rik/Ncbp2as2 and tetraspanin 9, which are both decreased on APOE4 animals (supplementary data figure 1). The former is reduced in AD and the latter promotes alpha secretase activity in cultured cells [55-57].

In terms of PNN components in addition to HAPLN2, we observe that 6-sulfated CSPGs are reduced in human APOE4 expressing mice. Though our studies used Western blot so that we can’t be certain that sulfation is changed in PNNs in particular, C6 sulfation does render PNNs more susceptible to proteolysis and generally less inhibitory. It may also influence increase calcium sequestration by PNNs to influence the activity of voltage gated calcium channels at sites including the axon initial segment [58, 59]. Importantly, overexpression of the 6 sulfotransferase has been shown to improve cognition in aged animals [36].

Because brain lysates may include PNN components from multiple sources such as loose ECM and peri-nodes of Ranvier, we also stained fixed hippocampal tissue from human APOE3 and APOE4 expressing mice for perineuronal localized aggrecan. As compared to WFA positive nets, aggrecan positive nets are relatively more abundant in the hippocampus and aggrecan immunostaining is also more abundant in aged mice [48]. Our results showed an APOE4 associated increase in hippocampal aggrecan intensity. Caveats of our aggrecan immunostaining studies, however, include a relatively low n and the sole use of females.

In the hippocampus, PNNs predominantly surround PV expressing inhibitory neurons. In the CA2 region however, they also surround excitatory neurons [60]. Changes in hippocampal PNNs have implications for cognitive flexibility, which may be impaired by excess PNN deposition, as well as oscillatory activity that can influence mood or working memory [21]. Of relevance is recent work that showed increased interneuron excitability in the APP/PS1 murine model of AD [61, 62], in that PV associated PNNs can increase interneuron excitability [20]. A recent study also showed increased levels of PNN components including tenascin and brevican in the APP/PS1 murine model of AD [63]. In contrast, PNN attenuation has been associated with amyloid plaque burden in the 5XFAD model [64]. Future studies could further explore that contribution of PV-localized PNN changes to interneuron excitability in amyloid depositing and/or APOE4 expressing mice.

Overall, the present study adds to our understanding of ECM dysregulation in conditions that increase AD risk. Future studies are warranted to determine whether excess ECM remodeling and deposition contribute to AD associated deficits in mood or cognition, in which case adjunct therapeutics to reduce excess ECM remodeling could be considered for high-risk individuals.

## Abbreviations

PNN: perineuronal net
HAPLN2: hyaluronan and proteoglycan link protein 2
HTRA1: high temperature requirement peptidase-1
CSPG: chondroitin sulfate proteoglycan
ECM: extracellular matrix
APOE: apolipoprotein E

## Figure Legends

***Supplemental Data Figure 1***.

Volcano plots proteomics results from mouse cohorts used in the study. HAPLN2 was among the most elevated proteins in the APOE4 compared to APOE3 mice. Additional proteins of interest to AD are highlighted in pink or green. A full excel file of significantly changed proteins by the analysis described herein, and a list of all proteins detected, is available on request via the corresponding author

## Acknowledgements

We very much appreciate National Institutes of Health training grant and R01 funding as follows: T32AG071745 (MA), T32NS041218 (ZC), and AG077002 (KC). We also thank Dr. Junfeng Ma and the Georgetown University Medical Center Proteomics Core Facility.

